# OPTIMIZED NEURON TRACING USING *POST HOC* REANALYSIS

**DOI:** 10.1101/2022.10.10.511642

**Authors:** Sara Azzouz, Logan A Walker, Alexandra Doerner, Kellie L. Geisel, Arianna K. Rodríguez Rivera, Ye Li, Douglas H Roossien, Dawen Cai

## Abstract

Over the last decade, the advances in Brainbow labeling allowed labeling hundreds of neurons with distinct colors in the same field of view of a brain [1, 2]. Reconstruction (or “tracing”) of the 3D structures of these images has been enabled by a growing set of software tools for automatic and manual annotation. It is common, however, to have errors introduced by heuristics used by tracing software, namely that they assume the “best” path is the highest intensity one, a more pertinent issue when dealing with multicolor microscope images. Here, we report *nCorrect*, an algorithm for correcting this error by reanalyzing previously created neuron traces to produce more physiologically-relevant ones. Specifically, we use a four dimensional minimization algorithm to identify a more-optimal reconstruction of the image, allowing us to better take advantage of existing manual tracing results. We define a new metric (hyperspectral cosine similarity) for describing the similarity of different neuron colors to each other. Our code is available in an open source license and forms the basis for future improved neuron tracing software.

## 1. INTRODUCTION

Advances in microscopy methods have recently enabled the high-throughput reconstruction of hundreds of neurons’ 3D structure in a single experiment. Recent studies, such as those by the BRAIN initiative [4], have demonstrated the value of large numbers of neuron reconstruction into understanding the structure of the nervous system through identifying roles of individual cell types. In order to process these data, large numbers of manual and automatic analysis applications have been developed [3, 5, 6, 7]. Despite these technical advances, a major limiting factor remains the proofreading and data analysis of neuron reconstructions which must be performed [8]. Proofreading includes verifying that neuron traces accurately represent the physiology of the target neuron, and are not overfit to the boundary of the neuron, which can occur when following the maximum intensity of the data (Fig. 1A).

**Fig. 1.**
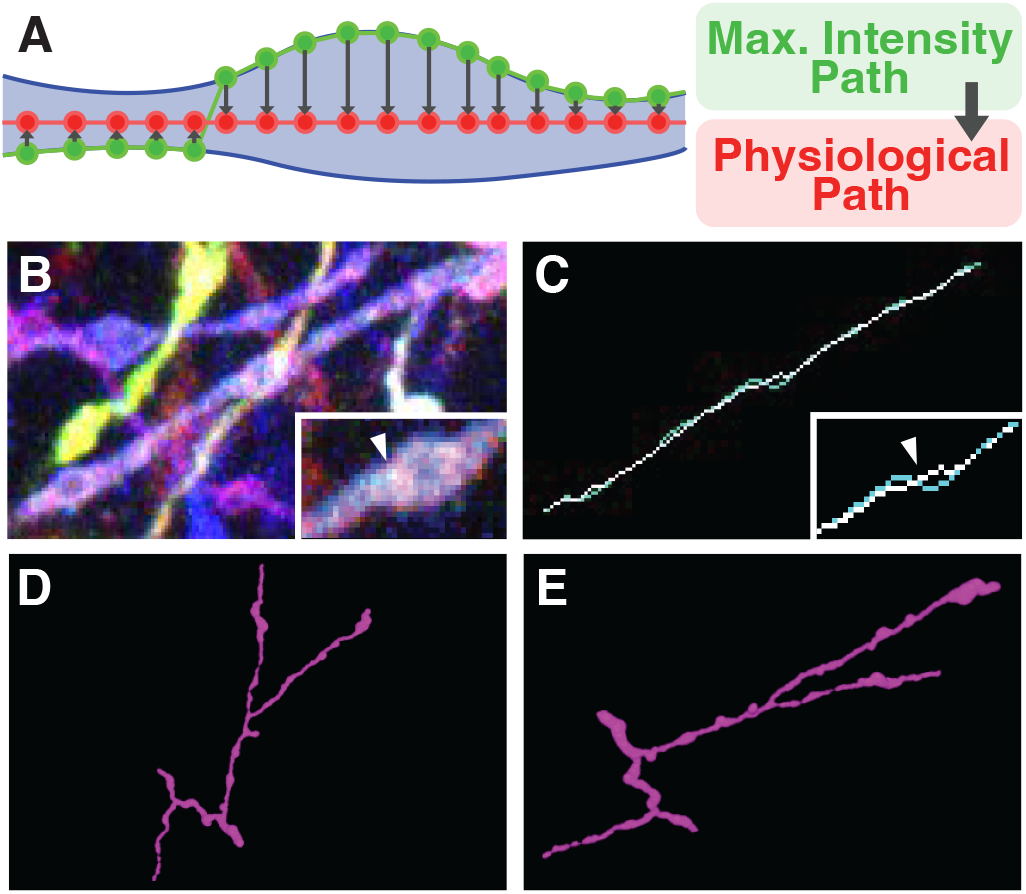
**A**, A schematic of the expected result of *nCorrect*. **B-C**, The result of applying *nCorrect* to data from [3]. Notably, *nCorrect* fixes a misplaced trace in a thick neurite. In **C**, white represents *nCorrect* result, while cyan represents the original trace. **D-E**, Two 3D views of reconstructed neurites, demonstrating the ability to estimate the volume of a neurite.

In our work, we have found many erred neuron reconstructions where the path is placed along the membrane edge of a neurite, due to that being the highest intensity point in the region of the image (e.g., Fig. 1B). Multi-color labeling techniques, such as Brainbow [1] or Tetbow [9], complicate this further becasue multiple neurons are often imaged within the same sample in different combinations of color, leading to the possibility that multiple neurons are in close proximity to each other. To ensure the fidelity of these reconstructions, we created *nCorrect*, an algorithm which uses a novel metric *multispectral cosine similarity* and a iterative minimization approach to identify optimized neuron traces in multispectral images.

## 2. METHODS

### 2.1. Algorithm motivation

*nCorrect* is motivated by an observation that many existing tools do not accurately model neurons. Specifically, because neurons are 3D objects, one must fit both a 3D segmentation as well as a skeleton to the raw data. Recently, the *SNT* [5] Fiji plugin [10] added a utility to perform this type of fitting, however it relies upon fitting of cross sections and using a thresholded monochrome image, which is incompatible with hyperspectral data without modification. Instead, we directly fit 3D spheres to the data, using a collection of techniques to account for the complexity of multispectral data, as described below.

### 2.2. Representation

We define a node *n* as a coordinate tuple with three integer spatial coordinates (*n*_*x*_, *n*_*y*_, *n*_*z*_) ∈*I* and a radius *n*_*r*_ ∈ ℤ ∩ [1, *n*_*r*,max_], where *I* represents a grid of all pixel values in a given micrograph image made up of a number of color channels *k*. Thus, each node represents a sphere in ℝ^3^. Finally, each node contains a vector of intensities 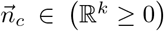. Neuron traces are represented as directed, acyclic chains or “paths” of nodes which are bounded in *I*.

### 2.3. Hyperspectral cosine similarity

To handle multispectral images (ones captured with multiple colors of light), our heuristic function uses a distance function based on the angle between different colors of light. Assuming that the color composition of each neuron is relatively uniform, we define a neuron’s reference color vector as the vector average of color over a path 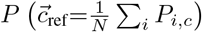. Using this, we can define the cosine similarity between any normalized color vectors 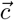 and 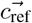 using the dot product:

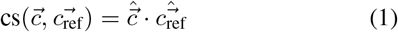

Here, we define a node’s color background ratio *n*_bg,col_ as the percentage of pixels which meet the criteria 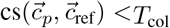.

### 2.4. Intensity thresholding

To identify areas of oversegmentation (background being included in the segmentation), we define a node’s background ratio *n*_bg,int_ as the ratio of pixels where the sum of a pixel’s color vector components 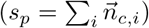 and a the sum of 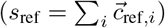 fail to satisfy *T* ^−^ *< s*_*p*_*/s*_ref_ *< T* ^+^. Here, the lower threshold (*T* ^−^) is defined by the function:

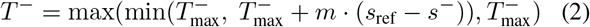

where 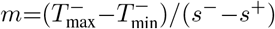 and parameters used are defined in Table 1. By setting *T* ^−^ as a function of *s*_ref_, we can better ensure that sections of low-intensity neurons are still classified as foreground while dim regions are not included as foreground for high-intensity neurons. Finally, after *n*_bg,int_ is computed, we can classify the node as a background node if *n*_bg,int_ *> T*_bg,int_. For background nodes where *n*_*r*_ *>* 1, we rectify *n*_bg,int_ to a value of 100 to disadvantage this node in the pathfinding procedure.

**Table 1.**
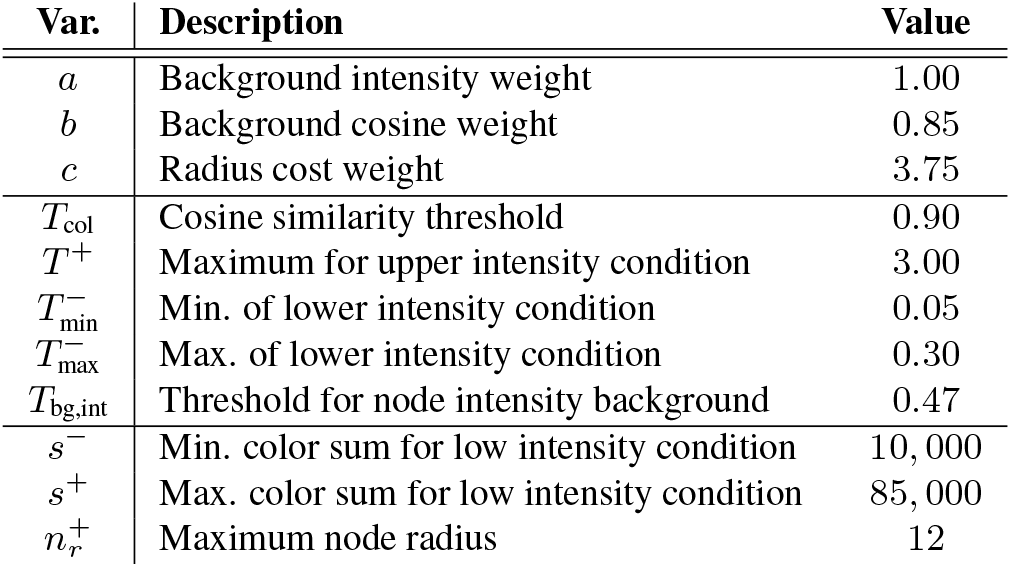
Parameters and the values used in this paper. Each parameter was determined by iterative searches executed in batch and may vary depending on the nature of the image being analyzed.

### 2.5. Cost function

Using *n*_bg,col_, *n*_bg,int_ as defined above, we define the heuristic cost function of node *n* as follows:

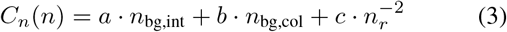

Where *a, b*, and *c* are given in Table 1. Thus the total cost function of a path can be computed using a sum across all nodes’s cost (*C*(*P*)= Σ_*i*_ *C*_*n*_ (*P*_*i*_)).

### 2.6. Iterative minimization approach

In order to obtain a corrected path along a neuron, we propose an iterative minimization approach (Algorithm 1), seeking to minimize the global penalty *C*(*P*). We begin by assuming that we have an *a priori* path, *P* ^0^ which is given as input to our algorithm and may represent the output from another tracing software. The radius of each node in *P* ^0^ is set by finding 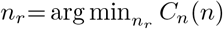. Next, we iterate through all nodes excluding the endpoints, calculating the penalty at each neighboring coordinate-radius pair. We then swap the current node with the most optimal neighboring node if it has a lower penalty value. This procedure is repeated until the algorithm converges to a minimal penalty path (i.e., no swaps are made over an entire iteration). Note that this iterative minimization approach is separately performed on each branch in a neuron, and gaps between branches and sub-branches are closed with a previously-described [3] A* search strategy before subsequent iterative minimization of sub-branches.

#### Algorithm 1 The iterative minimization algorithm

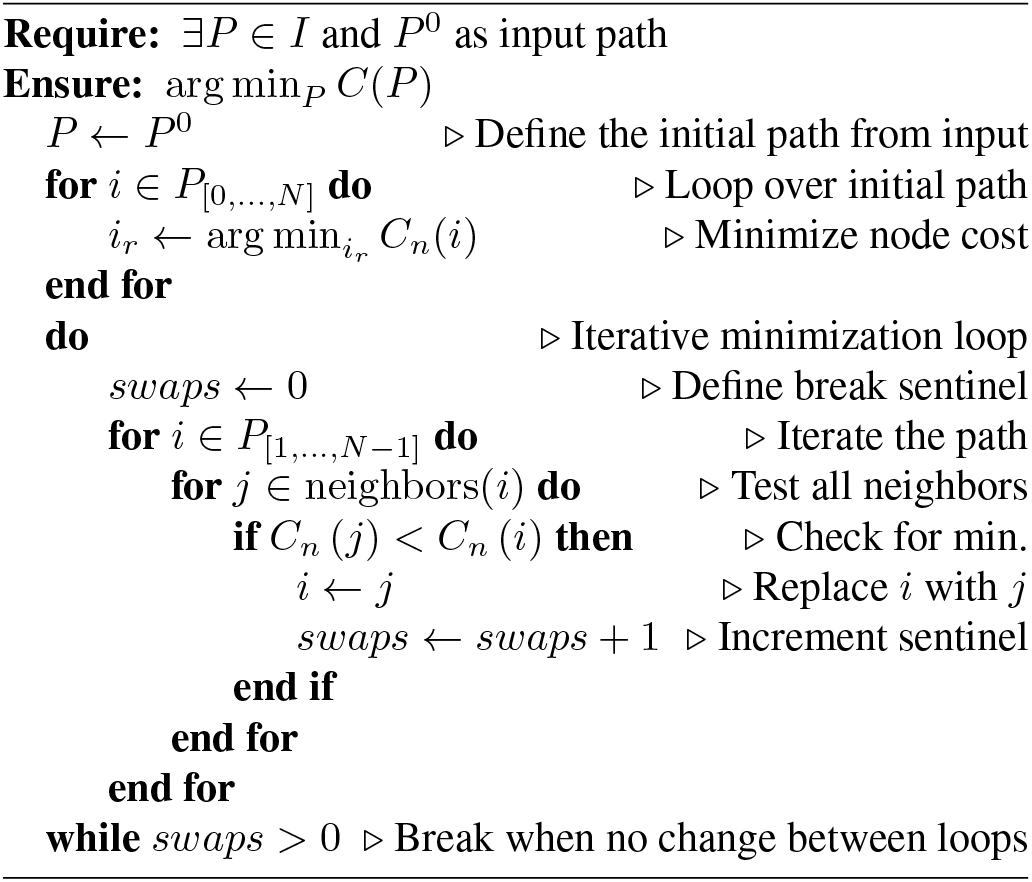

### 2.7. Code implementation

*nCorrect* was written in Java to ensure cross-platform portability, and to work well with the popular ImageJ/Fiji [10] image analysis environment for file loading and memory management. We have released this tool as a command line utility that can be run in batch scripts.

### 2.8. Biological sample preparation and analysis

PV-Cre (Pvalb^tm1(cre)Arbr^; Jackson Laboratory #008069) mice were injected into V1, immunolabeled, processed, and imaged as described previously [11]. Manual tracing of Brainbow data sets was done using *nTracer* 1.4 [3]. Each individual SWC file was exported from these analyses to be used as *nCorrect* inputs.

## 3. RESULTS

### 3.1. Cosine similarity and thresholding approach successfully segments neurons

To demonstrate the novel cosine similarity metric’s ability to segment neurons, we applied it to previously published data from [3] (Fig. 2A). After choosing a reference point (white ‘X’ in Fig 2A), the cosine similarity was calculated for each pixel using *n*_*r*_=2 (Fig. 2B). Each pixel was then thresholded for cosine similarity *<* 0.9 and intensity *<* 20% of maximum range (Fig. 2C-D). This demonstrates that our thresholding approach can be used to segment a neuron from both other neurons and from background.

**Fig. 2.**
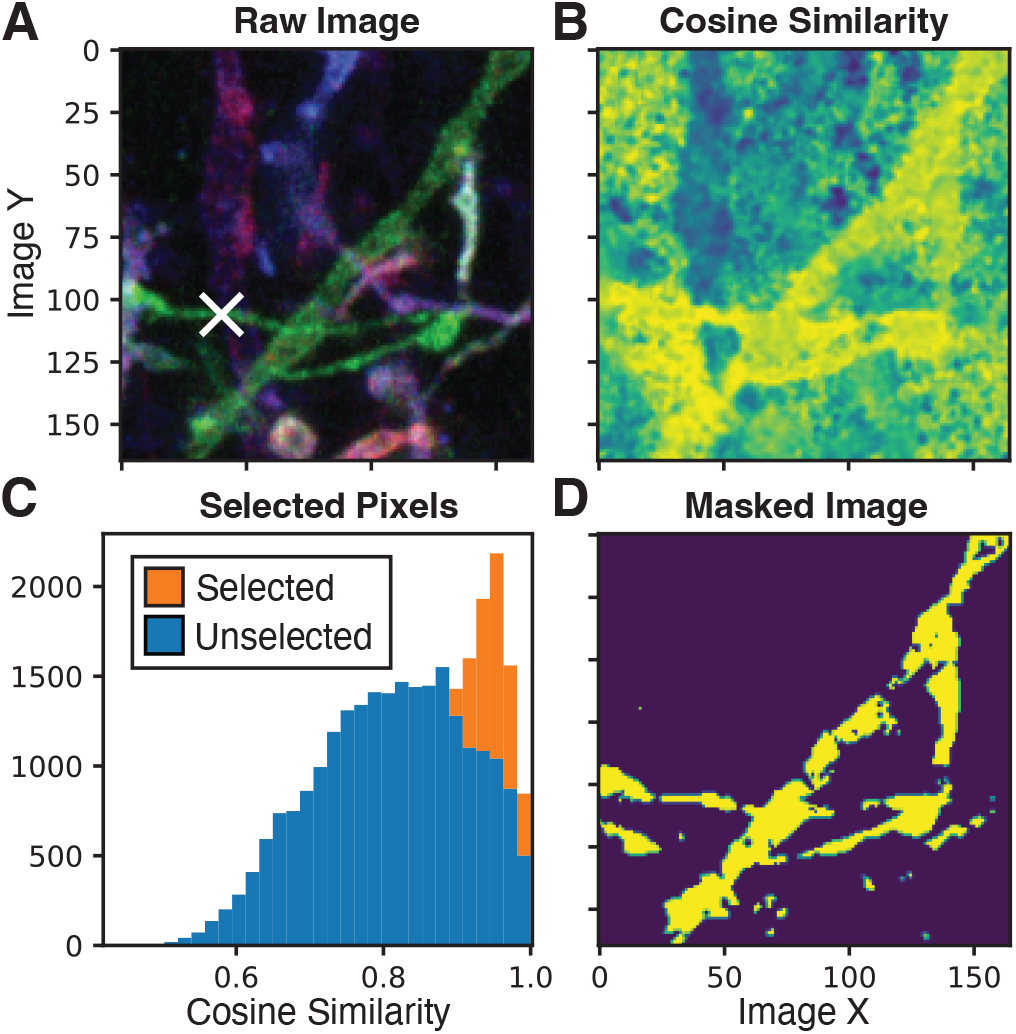
Use of cosine similarity scores for neuron discrimination. **A** A 20 frame 2D z-projection of an example image. The original image is 4 spectral channels, which are plotted as RGBR. A white ‘X’ represents a chosen sample point to select the green neurite in the image. **B** The cosine similarity score for each pixel is plotted against the reference point in **A. C** A distribution of all pixels in the image, separated by if a pixel meets set cosine similarity and intensity thresholds. **D** A plot of the pixels in **C**, demonstrating that the green neuron has been segmented by the cosine similarity metric.

### 3.2. *nCorrect* may result in neuron morphometrics changes

To evaluate the effect that correcting, we calculated a collection of neuron morphometrics for a collection of 70 individual tracing results with *nGauge* [12]. Three morphometrics were calculated, including the total length of the traced neurite, the total volume of the traced structure, and neuron tortuosity (Fig. 3). We find that path length and tortuosity in particular have a wide range of changes, which we attribute to path smoothing as a result *nCorrect* processing.

**Fig. 3.**
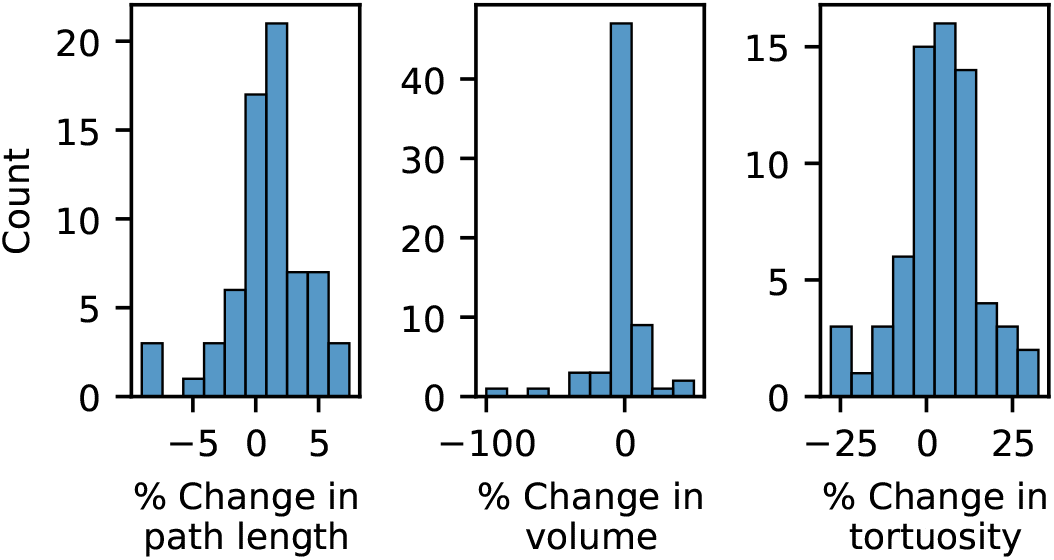
Morphometrics change in response to *nCorrect* processing. For each compared traced neurite, total path length, volume, and tortuosity was calculated for the original neurite and *nCorrect* results, and compared by 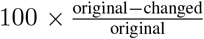. For each comparison, metrics with a value of zero were excluded from analysis because they have an indeterminate percentage change.

### 3.3. Human evaluation of *nCorrect* results

To evaluate the fidelity of *nCorrect*’s tracing results, 70 individual neurite tracing results were exported as individual SWC [13] neuron structure files for modification with *nCorrect*. The original and *nCorrect* results were assigned either a cyan or magenta color at random and distributed to four blind evaluators with moderate to advanced tracing experience. For each pair of results, evaluators were asked which trace displayed a more accurate trace of the neurite in the raw image, with the following parameters prioritized: maintains correct neurite (based on color), trace centered in the neurite, and trace smoothness. These evaluations were labeled “Improved” (*nCorrect* chosen), “Neutral” (neither was chosen), or “Error” (original result was chosen). For tracing results in the “Error” category, one expert evaluator further compared original and *nCorrect* tracing results to determine if the error was due to an error in the original *nTracer* result (Fig. 4B-C) or introduced by the *nCorrect* modification (Fig. 4D-E). Proportions of each category were averaged together across the four evaluators.

**Fig. 4.**
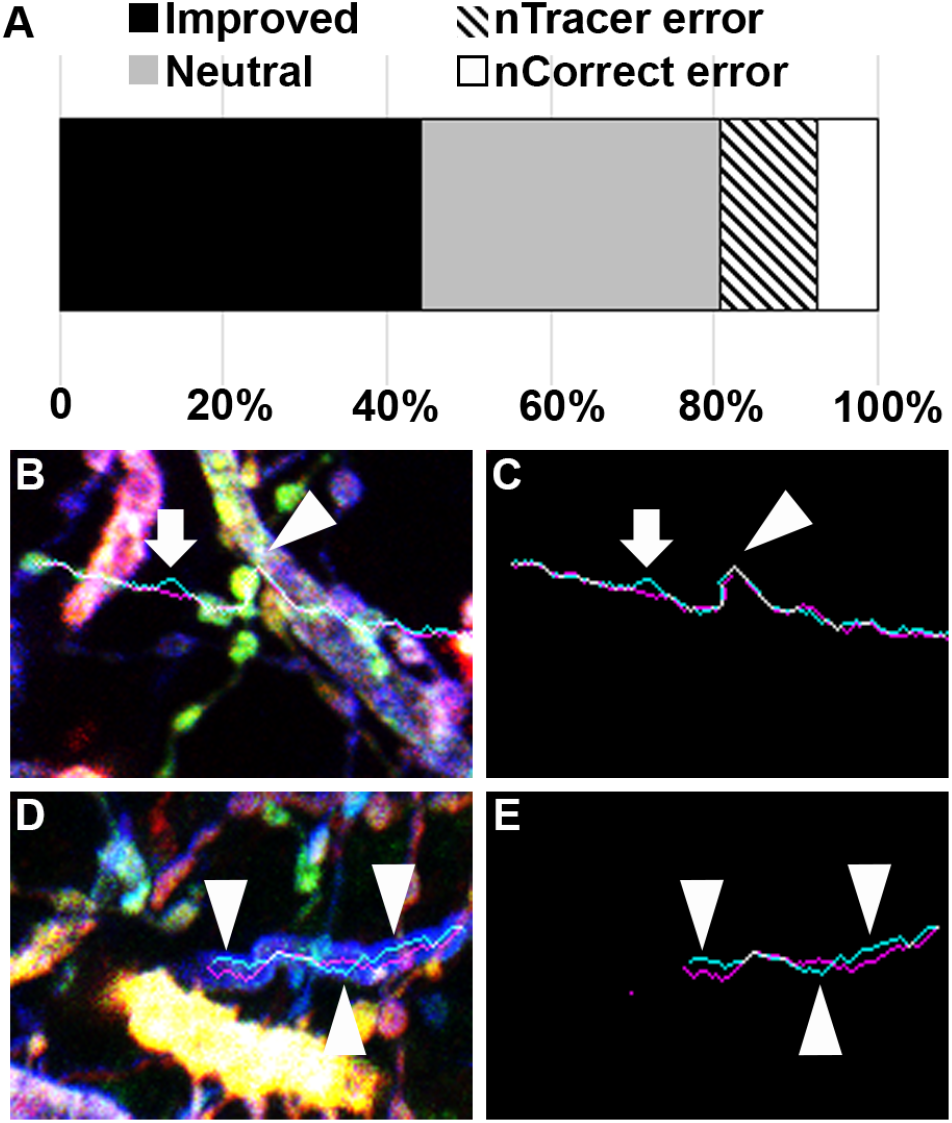
Human evaluation of nCorrect tracing fidelity. **A**, Proportions of evaluation results averaged from three blind evaluators, indicating that the vast majority (80.8%) of inputs were found to either by improved or unaffected by the *nCorrect* protocol. **B-C**, An example of an *nTracer* (magenta) error causing an error in the *nCorrect* (cyan) result. The *nTracer* result error is indicated by arrowhead, where the trace jumps to the wrong neurite before navigating back to the original neurite. The erroneous color sampling from the wrong neurite causes a downstream *nCorrect* error indicated by the arrow. **D-E** An example of an *nCorrect* error. In this example, the *nCorrect* result (cyan) prefers to hug the edge of the neurite rather than traverse the center of the neurite (indicated by the arrowheads in **D-E**). The *nTracer* result is shown in magenta.

From this blinded qualitative evaluation, the evaluators found that 44.1% of tracing results showed the *nCorrect* results were an improvement over the original results and that 36.7% were similar to the original results. For the remaining 19.2%, a *post-hoc* evaluation of traces determined that the source of errors in the majority of *nCorrect* tracing results were due to errors in the original *nTracer* result (Fig. 4B-C). Most of these occurred in highly dense regions where the *nTracer* trace jumped to the incorrect neurite (based on color composition), which either caused the *nCorrect* trace to follow to the incorrect neurite or, as shown in Fig. 4B-C, caused a nCorrect pathfinding error downstream of the *nTracer* error.

## 4. DISCUSSION AND CONCLUSION

*nCorrect* is the first algorithm to our knowledge, designed to improve neuron tracing fidelity in multispectral data that has already been reconstructed. We believe that this is a crucial step to large-scale neuron reconstruction protocols, as it not only gives a more accurate reconstruction, but also provides information about neuron width, etc. which are not present in many neuron reconstruction outputs currently.

We believe that versions of *nCorrect* can be used to further improve existing tracing software by optimizing the algorithm to run as a real-time step of current analyses. This could be accelerated by a C++ implementation, which could be significantly faster than the current Java implementation. Further, because this algorithm can be parallelized by neuron branch, it may be possible to accelerate it using GPU computing techniques.

## 5. CODE AVAILABILITY

A reference implementation of *nCorrect* is available from our lab’s [GitHub page](https://github.com/Cai-Lab-at-University-of-Michigan/nCorrect).

## 6. COMPLIANCE WITH ETHICAL STANDARDS

This study was performed in line with the principles of the Declaration of Helsinki. Approval was granted by the Institutional Review Board of the University of Michigan (PRO9289).

## 7. ACKNOWLEDGMENTS

LAW received support from a University of Michigan Rackham Graduate School Predoctoral Fellowship. This work was supported by NSF-1707316 (Neuronex-MINT), NIH-RF1MH123402, and NIH-RF1MH124611. The authors thank Chang Ge and Bin Duan for their comments on mathematical notation.

## 8. AUTHOR CONTRIBUTIONS

DC and LAW conceptualized *nCorrect*, which was then implemented by SA. All authors participated in neuron annotation experiments and software testing. DR generated the mouse microscopy images used for testing, and coordinated evaluation efforts. The manuscript was written by LAW, SA, DR, and DC, with edits and approval by all authors.

## REFERENCES

[1] Jean Livet, Tamily A Weissman, Hyuno Kang, Ryan W Draft, Ju Lu, Robyn A Bennis, Joshua R Sanes, and Jeff W Lichtman, “Transgenic strategies for combinatorial expression of fluorescent proteins in the nervous system,” Nature, vol. 450, no. 7166, pp. 56–62, Nov. 2007.

[2] Dawen Cai, Kimberly B Cohen, Tuanlian Luo, Jeff W Lichtman, and Joshua R Sanes, “Improved tools for the brainbow toolbox,” Nat. Methods, vol. 10, no. 6, pp. 540–547, May 2013.

[3] Douglas H Roossien, Benjamin V Sadis, Yan Yan, John M Webb, Lia Y Min, Aslan S Dizaji, Luke J Bogart, Cristina Mazuski, Robert S Huth, Johanna S Stecher, Sriakhila Akula, Fred Shen, Ye Li, Tingxin Xiao, Madeleine Vandenbrink, Jeff W Lichtman, Takao K Hensch, Erik D Herzog, and Dawen Cai, “Multispectral tracing in densely labeled mouse brain with ntracer,” Bioinformatics, vol. 35, no. 18, pp. 3544–3546, Sept. 2019.

[4] BRAIN Initiative Cell Census Network (BICCN), “A multimodal cell census and atlas of the mammalian primary motor cortex,” Nature, vol. 598, no. 7879, pp. 86–102, Oct. 2021.

[5] Cameron Arshadi, Ulrik Günther, Mark Eddison, Kyle I S Harrington, and Tiago A Ferreira, “SNT: a unifying toolbox for quantification of neuronal anatomy,” Nat. Methods, vol. 18, no. 4, pp. 374–377, Apr. 2021.

[6] Yimin Wang, Qi Li, Lijuan Liu, Zhi Zhou, Zongcai Ruan, Lingsheng Kong, Yaoyao Li, Yun Wang, Ning Zhong, Renjie Chai, Xiangfeng Luo, Yike Guo, Michael Hawrylycz, Qingming Luo, Zhongze Gu, Wei Xie, Hongkui Zeng, and Hanchuan Peng, “TeraVR empowers precise reconstruction of complete 3-D neuronal morphology in the whole brain,” Nat. Commun., vol. 10, no. 1, pp. 1–9, Aug. 2019.

[7] Ludovica Acciai, Paolo Soda, and Giulio Iannello, “Automated neuron tracing methods: An updated account,” Neuroinformatics, vol. 14, no. 4, pp. 353–367, Oct. 2016.

[8] Hanchuan Peng, Fuhui Long, Ting Zhao, and Eugene Myers, “Proof-editing is the bottleneck of 3D neuron reconstruction: the problem and solutions,” Neuroinformatics, vol. 9, no. 2-3, pp. 103–105, Sept. 2011.

[9] Richi Sakaguchi, Marcus N Leiwe, and Takeshi Imai, “Bright multicolor labeling of neuronal circuits with fluorescent proteins and chemical tags,” Elife, vol. 7, Nov. 2018.

[10] Johannes Schindelin, Ignacio Arganda-Carreras, Erwin Frise, Verena Kaynig, Mark Longair, Tobias Pietzsch, Stephan Preibisch, Curtis Rueden, Stephan Saalfeld, Benjamin Schmid, Jean-Yves Tinevez, Daniel James White, Volker Hartenstein, Kevin Eliceiri, Pavel Tomancak, and Albert Cardona, “Fiji: an open-source platform for biological-image analysis,” Nat. Methods, vol. 9, no. 7, pp. 676–682, June 2012.

[11] Douglas H Roossien and Dawen Cai, “Imaging neural architecture in brainbow samples,” Methods Mol. Biol., vol. 1642, pp. 211–228, 2017.

[12] Logan A Walker, Jennifer S Williams, Ye Li, Douglas H Roossien, Wei Jie Lee, Nigel S Michki, and Dawen Cai, “ngauge: Integrated and extensible neuron morphology analysis in python,” Neuroinformatics, Mar. 2022.

[13] Sumit Nanda, Hanbo Chen, Ravi Das, Shatabdi Bhattacharjee, Hermann Cuntz, Benjamin Torben-Nielsen, Hanchuan Peng, Daniel N Cox, Erik De Schutter, and Giorgio A Ascoli, “Design and implementation of multi-signal and time-varying neural reconstructions,” Sci Data, vol. 5, pp. 170207, Jan. 2018.

